# Heritability of skewed X-inactivation in female twins is tissue-specific and dependent on age

**DOI:** 10.1101/593251

**Authors:** Antonino Zito, Matthew N. Davies, Pei-Chien Tsai, Susanna Roberts, Stefano Nardone, Jordana T. Bell, Chloe C. Y. Wong, Kerrin S. Small

**Affiliations:** Department of Twin Research & Genetic Epidemiology, King’s College London, London SE1 7EH, UK; The Francis Crick Institute, 1 Midland Road, London NW1 1AT, UK; Institute of Psychiatry, Psychology & Neuroscience, King’s College London, London SE5 8AF, UK; Division of Endocrinology, Diabetes, and Metabolism, Department of Medicine, Beth Israel Deaconess Medical Center, Harvard Medical School, Boston, MA 02215, USA

## Abstract

To balance the X-linked transcriptional dosages between the sexes, one of the two X-chromosomes is randomly selected to be inactivated in somatic tissues of female placental mammals. Non-random, or skewed X-chromosome inactivation (XCI) toward one parental X has been observed in female somatic tissues, and this skewing effect has been associated with several complex human traits. However, the extent of the influence of genetic and environmental factors on XCI skewing is largely unknown. Here, we use RNA-seq and DNA-seq data taken from a large cohort of female twins to quantify the degree of skewing of XCI (DS) in multiple tissues and to study the relationship of XCI with age, genetic factors and complex traits. We show that the XCI patterns are highly tissue-specific with a higher prevalence of skewed XCI in blood-derived tissues than in fat or skin tissue. We also show that the DS in blood-derived tissues is associated with age and that the acquired DS occurs uniquely in blood-derived tissues with an inflection point at approximately 55 years of age. Heritability analysis indicates that the heritability of DS is both age and tissue specific; DS is heritable in blood tissue of females >55 years-old (h^2^ = 0.34) but is not heritable in blood tissues of females <55 years-old (h^2^ = 0), nor in skin and fat tissues at any age. We find a positive association between the DS and smoking status in blood tissues of older females (*P* = 0.02). The high tissue specificity of XCI patterns in human indicates the existence of tissue-specific mechanisms influencing XCI patterns, including genetic and environmental factors. We conclude that the heritability of XCI skewing in blood-derived tissues is dependent on age, representing a Gene x Age interaction that can shift the functional allelic dosage of an entire chromosome in a tissue-restricted manner.

## Introduction

To balance the X-linked transcriptional dosages between the single X chromosome of males and the two X chromosomes of females, one X chromosome is silenced in female placental mammals^1^. The X-chromosome inactivation (XCI) process starts during preimplantation phases of human embryonic development, presumably at around the 8-cell stage^2^. XCI is initiated by the transcription of *XIST*, a 17 kb, alternatively spliced long non-coding RNA mapped to Xq13.2 and exclusively expressed on the inactive X (Xi) ^3^. Once transcribed, *XIST* molecules spread in *cis* along the X chromosome^4,5^ inducing a progressive epigenetic silencing through the recruitment of chromatin remodelling enzymatic complexes, which impose repressive histone and DNA changes on the Xi chromosome^6,7^. Within each cell, the parental X chromosome selected for inactivation seems to occur at random, and the Xi is mitotically inherited to future somatic daughter cells. This random inactivation results in a mosaic of cells within an individual, where overall, a balanced expression (50:50) of both parental X-linked alleles is expected. Asymmetric selection of the X chromosome to inactivate causes the predominance of one parental Xi in a population of cells, unbalancing the X-linked transcriptional and allelic dosages toward one parental X chromosome. This phenomenon, known as skewed XCI (or non-random XCI), occurs when at least 80% of cells within a tissue inactivate the same parental X chromosome. The factors underlying primary skewed XCI are varied and several mechanisms are possible (reviewed in^8^). Secondary (or acquired) skewed XCI can result from positive selection of cells that after having inactivated a particular parental X, acquire a survival advantage over cells who inactivated the other parental X chromosome. Skewed XCI patterns can also be generated by the stochastic overrepresentation of cell clones in a given tissue, due for instance, to depletion of stem cell populations.

Comprised of 155 MB and containing >800 protein-coding genes, the X chromosome represents approximately 8% of the human genome. In heterozygous females with skewed XCI, the X-linked transcriptional and allelic dosages of silenced genes are unbalanced and may be functionally homozygous. Skewed XCI is a major cause of discontinuity of dominance and recessiveness, as well as penetrance and expressivity of X-linked traits. How skewed XCI patterns modulate phenotypes in females, and whether they are a cause or a consequence of associated phenotypes is not fully understood. Skewed XCI patterns have been observed in females with X-linked diseases^9,10^, autoimmune disorders^11,12^, as well as in breast^13^ and ovarian cancer^14^. In autoimmune diseases with higher prevalence in females, including rheumatoid arthritis and systemic lupus erythematosus, XCI is hypothesized to play a role. Chromosome X is enriched for immune-related genes and skewed XCI patterns cause the breakdown of thymic tolerance induction processes^15^ conferring a high predisposition to develop autoimmunity (reviewed in^16^). XCI skewing levels in blood-tissues have been associated with ageing, with multiple studies indicating an increase after 50-60 years of age^17-22^. To date, the mechanisms underlying skewed XCI in humans remain to be determined and hypotheses are still controversial. Several twin studies have reported that genetic factors contribute to XCI skewing in blood derived cells^21,23^, while other evidence indicated that most of the XCI skewing levels in human are acquired secondarily^24^.

Nearly all studies of XCI skewing levels in humans have been carried out in peripheral blood samples or in very small sample sizes^25^, while XCI patterns in other tissues have not been studied in great detail^19,26,27^. In this study, we comprehensively assessed XCI patterns in a multi-tissue sample of nearly 800 female twins from the TwinsUK cohort^28^. We quantified the degree of skewing of XCI using a metric based on *XIST* allele-specific expression from paired RNA-seq and DNA-seq data in four tissues. We examined the tissue-specific prevalence of skewed XCI patterns, compared the XCI skewing levels between tissues and evaluated the association between XCI skewing and lifestyle traits. In order to investigate the factors underlying the skewed XCI, we utilized classical twin models to characterize the extent of the influence of genetic and environmental factors on the tissue-specific skewed XCI, showing that the XCI patterns have both a heritable and environmental (age) basis.

## Results

### Quantification of degree of skewing in TwinsUK

We assessed XCI patterns in multi-tissue samples from female twin volunteers from the TwinsUK cohort aged 38–85 years old (median age = 60; Figure S1) ^28,29^. We quantified the degree of skewing of XCI using a metric based on *XIST* allele-specific expression (ASE) from paired RNA-seq and DNA-seq data. *XIST* is uniquely expressed from the Xi^3^, so the relative expression of parental alleles within the *XIST* transcript are representative of XCI skewing levels in a sample^27^. RNA-seq and genotype data were available for 814 Lymphoblastoid Cell Lines (LCL) samples, 395 whole-blood samples, 766 subcutaneous adipose tissue samples (herein referred as fat) and 716 skin samples. After stringent quality control, we obtained *XIST*_ASE_ calls for 422 LCL samples, 72 whole-blood samples, 378 fat samples and 336 skin samples. The smaller sample size for whole-blood was due to the relatively smaller size of the starting dataset and the relatively lower RNA-seq coverage of this tissue in our dataset. In order to have an absolute measure of the magnitude of the XCI skewing levels in each sample, we calculated the degree of skewing of XCI (DS) from the *XIST*_ASE_ calls. DS is defined as the absolute deviation of the *XIST*_ASE_ from 0.5 (see Materials and methods) and it has been similarly used in other investigations to assess XCI patterns^20,30,31^ and the XCI status of X-linked genes^32^. In line with previous investigations^21,33^ we classified samples with DS < 0.3 (corresponding to 0.2 < *XIST*_ASE_ < 0.8) to have random XCI, and samples with DS ≥ 0.3 (corresponding to *XIST*_ASE_ ≤ 0.2 or *XIST*_ASE_ ≥ 0.8) to have skewed XCI. Unless otherwise specified, DS was used in all the analyses performed.

We assessed the robustness of our estimates of the degree of skewing with an alternative DNA-based measure of XCI, the Human Androgen Receptor Assay (HUMARA)^34^. HUMARA was and is still the ‘gold standard’ technique to assess XCI patterns. A previous study has reported good replicability between HUMARA and expression-based quantification of XCI skewing^35^. We used HUMARA to measure XCI skewing levels in 10 archived whole-blood DNA samples obtained at the same clinical visit as the LCLs samples. Spearman’s correlation between the quantifications was 0.71, at *P* = 0.031, revealing a high degree of reproducibility between the both *XIST*_ASE_ and HUMARA methods and the LCLs and whole-blood (Fig 1a).

**Fig. 1:**
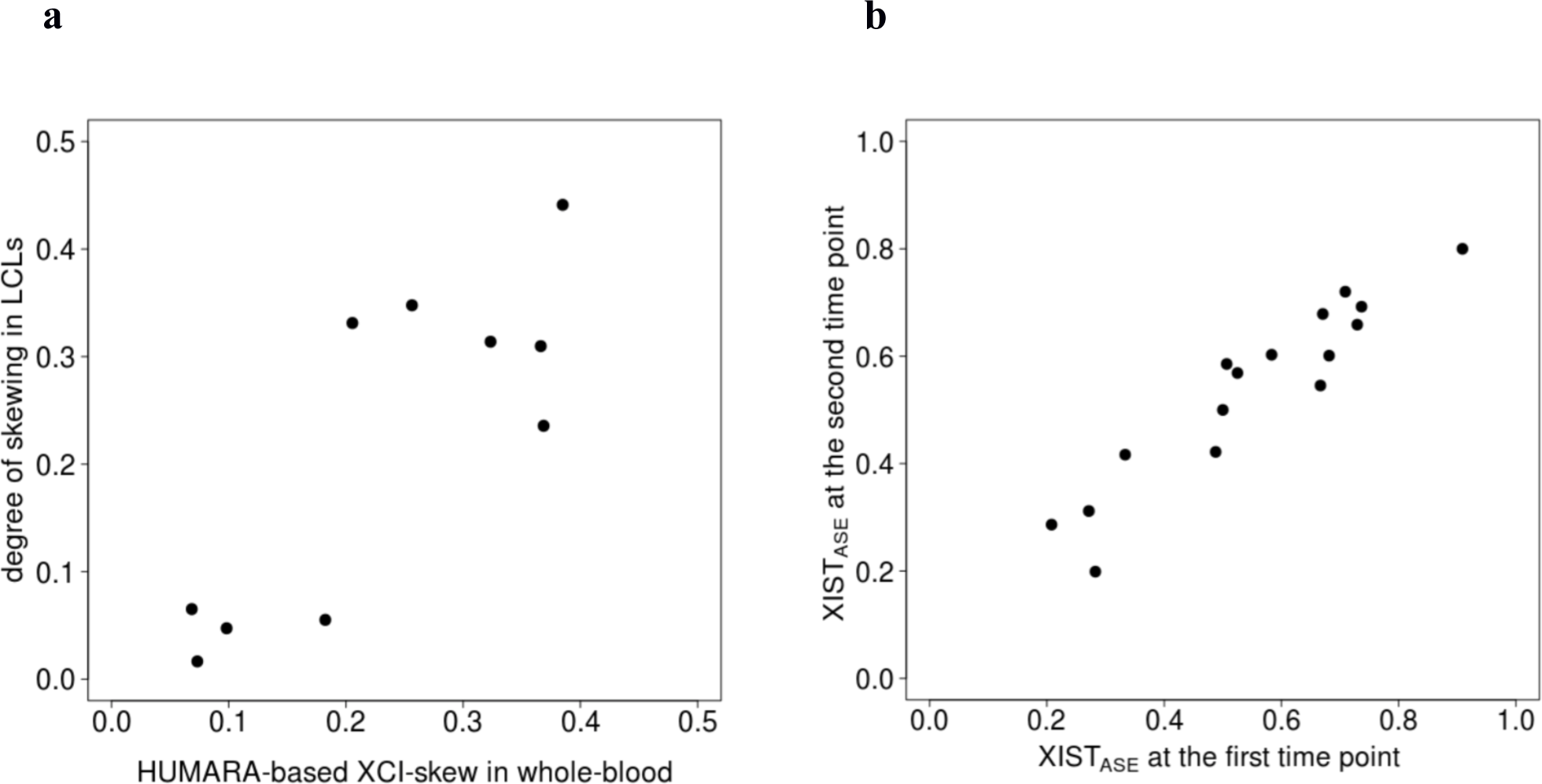
Comparison to HUMARA and longitudinal quantifications of XCI skewing levels reveal that *XIST*_ASE_ is a reproducible and accurate proxy of XCI skewing levels. **a** *Scatter plot* showing the comparison between the XCI skewing levels in 10 whole-blood and LCLs samples quantified with HUMARA and *XIST*_ASE_ respectively. **b** *Scatter plot* of the *XIST*_ASE_ at time point 1 and at time point 2 in 16 whole-blood samples.

Previous investigations have reported that the XCI skewing levels increase with age in blood tissues, as discussed below. While it would be expected that increases in XCI skewing levels would be observed over relatively large time spans, we would expect minimal variations of XCI skewing levels between 2 close time points. We therefore reasoned that the sensitivity of our quantifications could also be assessed by comparing the XCI skewing levels in the same individuals at close time points. Briefly, using a publicly-available longitudinal whole-blood RNA-seq dataset from the TwinsUK cohort^36^, we generated *XIST*_ASE_ calls at 2 time points (1 to 2.7 years later) in 16 samples (see the Materials and methods). Spearman’s correlation between the *XIST*_ASE_ calls at the first time point and the *XIST*_ASE_ calls at the second time point was 0.94 at *P* < 2 x 10^−15^ (Fig 1b). This indicates that *XIST*_ASE_ is a sensitive proxy when assessing the stability of XCI patterns over short time periods. Overall, these results indicate that *XIST*_ASE_ is a reproducible, accurate and sensitive proxy of XCI skewing levels.

### Skewed XCI is tissue-specific with higher prevalence in blood-derived tissues

We observed a wide range of DS values in the four tissues (Fig 2), with clear differences in the prevalence of skewed individuals between tissues. Blood-derived tissues had the highest incidence of skewed individuals, with skewed XCI observed in 34% of LCLs samples and 28% of whole-blood samples and a lower incidence in the primary tissues, where 12% of fat and 16% of skin samples exhibited skewed XCI (Table 1). In order to examine the extent of similarities of XCI patterns between tissues, we compared the tissue-specific XCI skewing levels in a pairwise manner (Fig 3). For each tissue-tissue comparison, we included individuals with *XIST*_ASE_ calls in both tissues (Table 2). We found the strongest correlation on *XIST*_ASE_ calls between LCLs and whole-blood (N = 59, r = 0.78, *P* = 2 × 10^-13^), indicating that blood-derived tissues share highly similar XCI skewing levels. We also found a good degree of similarity between the XCI skewing levels in fat and skin tissues (N = 252, r = 0.47, *P* = 2 × 10^-15^; Fig 3). However, low concordance was observed between skin and whole-blood (N = 47, r = 0.3, *P* = 0.04) and fat and whole-blood (N = 57, r = 0.33, *P* = 0.02). Our data demonstrate that tissue-specific XCI skewing within an individual is common in the population, indicating that XCI patterns are partially controlled by tissue-specific regulatory mechanisms.

**Table 1.** Prevalence of skewed X-inactivation (XCI) differs across tissues in the TwinsUK cohort.

| Tissue | Informative individuals | Skewed XCI individuals | Prevalence of skewed XCI |
| --- | --- | --- | --- |
| LCLs | 422 | 145 | 34% |
| Whole-blood | 72 | 20 | 28% |
| Fat | 378 | 45 | 12% |
| Skin | 336 | 54 | 16% |

**Table 2.** Correlation of XCI-skew varies between tissue pairs, and is highest between blood-derived tissues. Coefficient of correlation [and its p-value] of XCI-skew between tissue pairs.

| **Tissue** | **LCLs** | **Whole-blood** | **Fat** | **Skin** |
| --- | --- | --- | --- | --- |
| LCLs | * | 0.78 [2e-13] | 0.42 [5e-14] | 0.3 [1e-06] |
| Whole-blood | 0.78 [2e-13] | * | 0.33 [1e-02] | 0.3 [4e-02] |
| Fat | 0.42 [5e-14] | 0.33 [1e-02] | * | 0.47 [2e-15] |
| Skin | 0.3 [1e-06] | 0.3 [4e-02] | 0.47 [2e-15] | * |

**Fig. 2:**
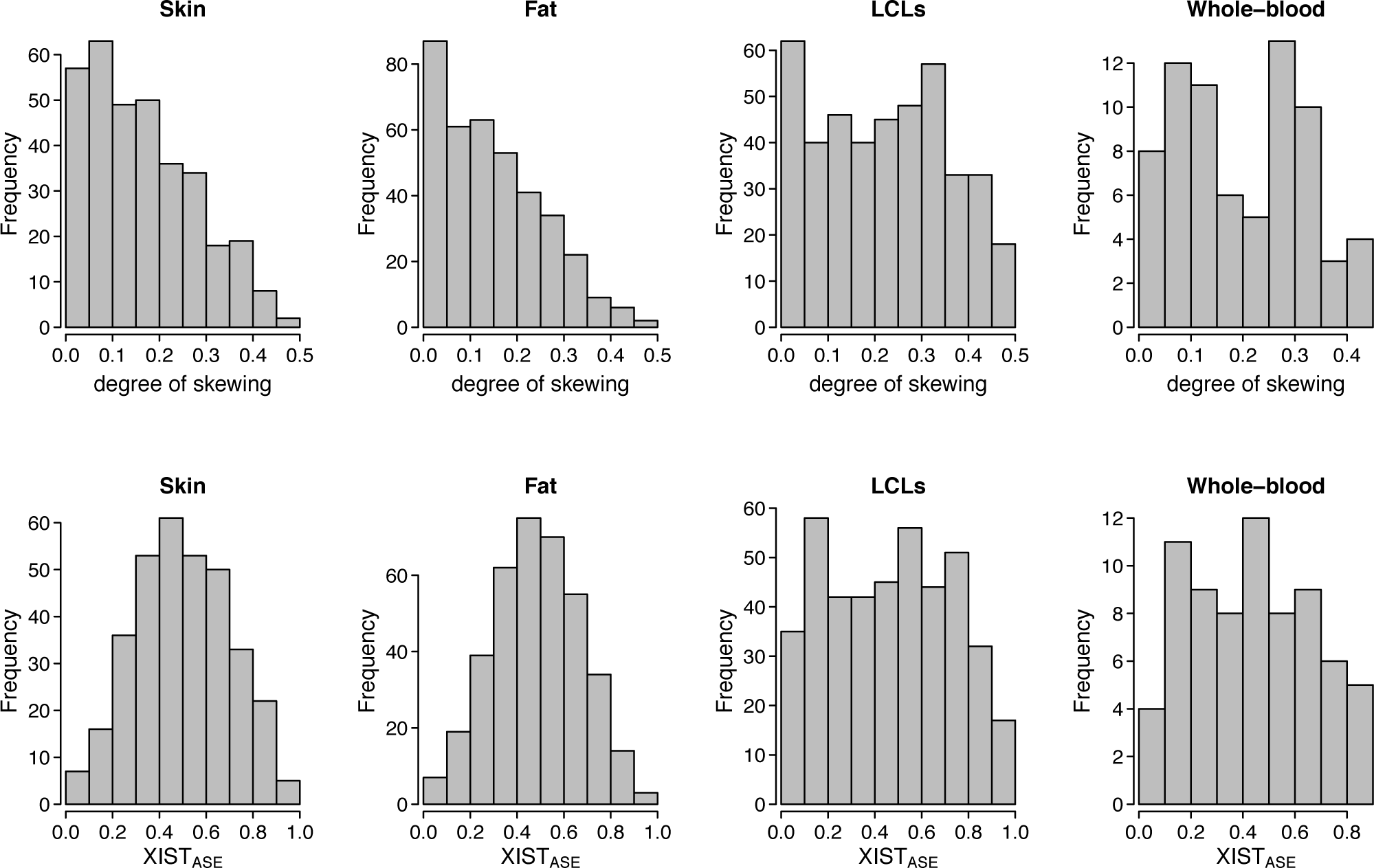
Skewed chromosome X inactivation varies across tissues with a higher prevalence of skewed samples in blood-derived tissues than fat and skin tissues. Distribution of the degree of skewing (top row) and *XIST*_ASE_ (bottom row) in LCLs (422 samples), whole-blood (72 samples), fat (378 samples) and skin (336 samples) tissues.

**Fig. 3:**
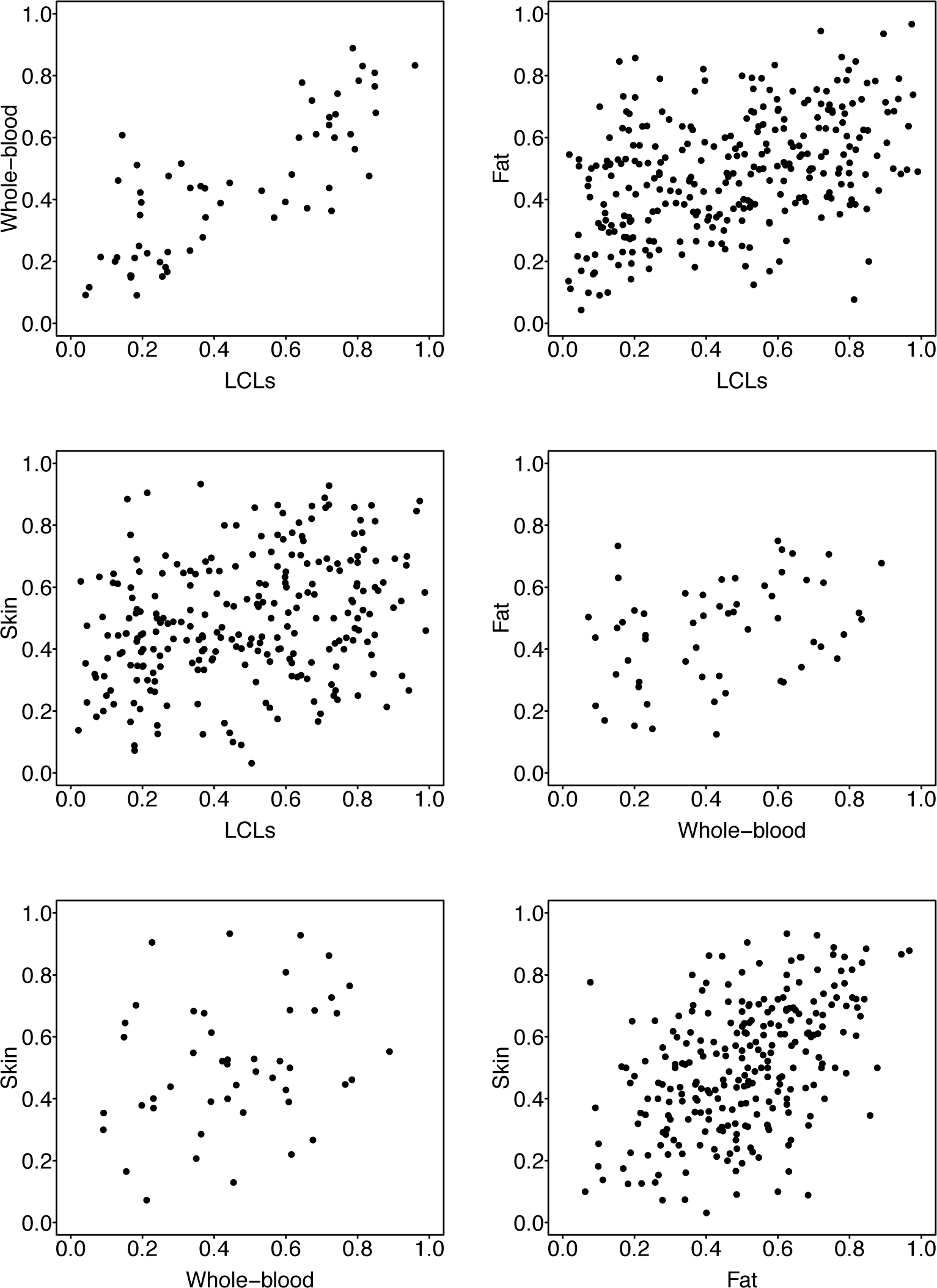
Pairwise tissue-tissue comparison of the *XIST*_ASE_ calls reveal highest similarity of XCI skewing levels between blood-derived tissues. Each plot shows the comparison of *XIST*_ASE_ between two tissues from an individual. Each dot represents an individual. In each comparison all individuals with available *XIST*_ASE_ calls for both tissues were included.

The active or inactive state of each X chromosome in a cell is clonally passed on to daughter cells. In a pool of cells derived from a single clone (or patch), the XCI patterns are expected to be completely skewed. Patch size refers to the amount of cell clones in a pool of cells (e.g. in a tissue biopsy). We considered the possibility that patch size might bias our quantification of XCI patterns in fat and skin samples. This is likely to occur in biopsies that are smaller than the tissue patch size. However, several considerations led us to exclude the possibility that patch sizes in fat and skin biopsies might confound our *XIST*_*ASE*_ calls. First, the biopsies included skin samples of 8*mm*^3^ in size, which were cut into 2 skin and 3 fat samples. As reported in another study, this size is large enough to measure the XCI ratio without being confounded by patch size^37^. Second, most individuals exhibit random XCI patterns in fat and skin tissues, which is unlikely if patch size was larger than the biopsies. We therefore conclude that the biopsies used in this study are large enough to accurately assess the XCI patterns without being biased by patch size.

### LCLs in this study are representative of XCI skewing *in-vivo* in blood tissues

Lymphoblastoid Cell Lines (LCLs) generated by Epstein-Barr virus mediated transformation of B lymphocyte cells have been and are widely used in gene expression studies. However, the possibility that the cell lines are monoclonal and/or polyclonal due to selection in the transformation process or clonal expansion in cell culture, and hence not be representative of the *in-vivo* XCI skewing levels, is a potential problem when using LCLs to assess XCI skewing^38^. As the profiled RNA in this study was extracted from the LCLs very shortly after transformation with limited passaging or time in culture we expected this effect to be minimal, however, to address the possibility we performed the following analyses. First, as described above and shown in Figure 1a, the degree of skewing in LCLs were highly correlated with the HUMARA-based quantifications of XCI patterns in paired whole-blood samples (r = 0.71, N = 10). We would not expect such high similarity between the two quantifications if clonal propagation had occurred in LCLs samples after preparation. This was confirmed by the high correlation between LCLs and whole-blood *XIST*_ASE_ values (r = 0.78, N = 59; Fig 3) and overall similarity in the prevalence of skewed XCI in LCLs and whole-blood (Table 1). Finally, we assessed the degree of skewing in monocytes, B, T-CD4^+^, T-CD8^+^ and natural-killer (NK) cells purified from two monozygotic twins exhibiting skewed XCI patterns in LCLs and from one individual exhibiting random XCI patterns in LCLs. We found that in both monozygotic twins showing skewed XCI in LCLs, the majority of immune cell types exhibited skewed XCI patterns. Conversely, none of the immune cell types purified from the non-skewed individual exhibited skewed XCI patterns (Table 3). We conclude that the XCI skewing levels of LCLs in this study are representative of XCI skewing *in-vivo* in blood tissues.

**Table 3.**
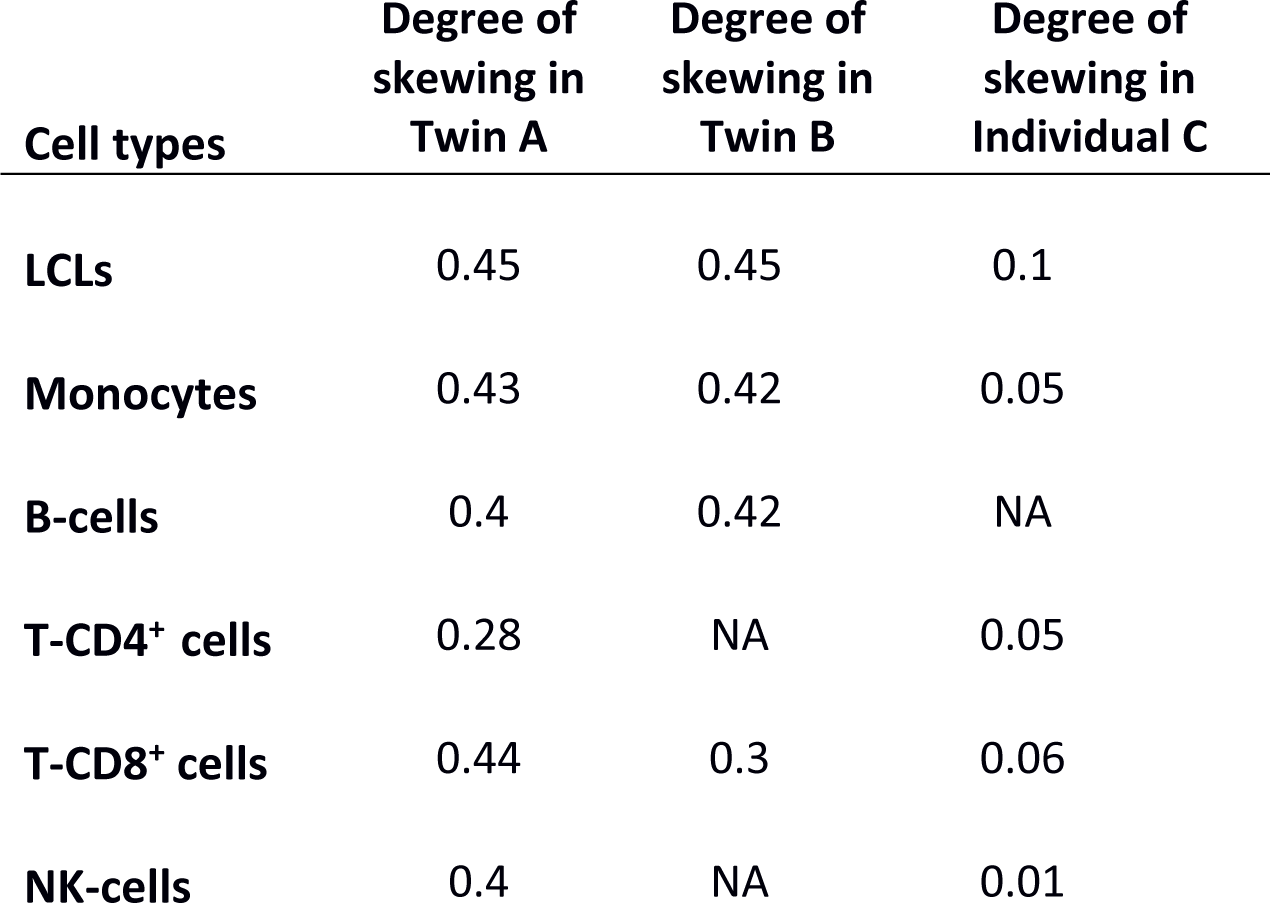
XCI-skew in LCLs in this study is representative of XCI in purified primary immune cells. Degree of skewing of XCI in immune cell types purified from 2 monozygotic twins (Twin A, Twin B) exhibiting skewed XCI patterns in LCLs, and from 1 individual (Individual C) exhibiting random XCI pattern in LCLs. Degree of skewing ≥ 0.3 indicates skewed XCI patterns.

| **Cell types** | **Degree of skewing in Twin A** | **Degree of skewing in Twin B** | **Degree of skewing in Individual C** |
| --- | --- | --- | --- |
| **LCLs** | 0.45 | 0.45 | 0.1 |
| **Monocytes** | 0.43 | 0.42 | 0.05 |
| **B-cells** | 0.4 | 0.42 | NA |
| **T-CD4<sup>+</sup> cells** | 0.28 | NA | 0.05 |
| **T-CD8<sup>+</sup> cells** | 0.44 | 0.3 | 0.06 |
| **NK-cells** | 0.4 | NA | 0.01 |

### XCI skewing levels are positively associated with age in blood-derived tissues

XCI skewing levels in peripheral blood have been shown to increase with age in multiple studies^17-21,23,35,39,40^. The age-related increase of XCI skewing levels continues throughout life, since centenarians exhibit higher XCI skewing levels than 95 years-old females^21^. However, there is very limited knowledge on the relationship between XCI patterns and ageing in tissues other than blood. In order to explore this, we investigated the association between age and degree of skewing in LCLs, fat and skin. Our whole-blood estimates were excluded from analysis due to low sample size (N = 72). Age was positively associated with XCI skew in LCLs (N = 422, *P* < 0.01), but we did not detect any association between XCI skew and age in skin (N = 336, *P* = 0.4) or in fat (N = 378, *P* = 0.7).

We next explored the dynamics of DS and age progression in each tissue, using the lowess procedure. Lowess curve detected an increase of DS beginning at around 55 years-old in LCLs (Fig 4), in agreement with what was found in other studies^19,21^. Since the increase of DS starts at around 55 years, we divided LCLs samples into a younger group (N = 141, age < 55) and an older group (N = 281, age ≥ 55). We found that the mean DS in LCLs was significantly higher in older than in younger females (DS_younger_ = 0.2, DS_older_ = 0.24, *P* = 0.03; Figure 4). Accordingly, we found that the frequency of skewed XCI in LCLs was significantly higher in older (38%) than in younger (28%) females (χ^2^ test, *P* = 0.04; Fig 4). In agreement with the lack of association between the DS and age, we did not detect significant differences between the mean DS in young and older females in fat (DS_younger_ = 0.15, DS_older_ = 0.15) or in skin tissues (DS_younger_ = 0.16, DS_older_ = 0.17). To acquire a more detailed view of the tissue-specific prevalence of skewed XCI in different groups of age, we categorized the samples into four age groups (40-50, 50-60, 60-70, >70) and calculated the frequency of skewed XCI in each category (Fig 4). We found that the frequency of skewed XCI increased with age in LCLs, with 41% of individuals >65 years-old demonstrating skewed XCI patterns. We did not observe any increase in the skewed XCI frequencies with age in fat and skin tissues. Overall, these data further confirm that XCI skewing levels increase with age in blood-derived tissues, supporting previous investigations. However, we find that there is no increase in XCI in fat and skin tissue from the same individuals, suggesting that acquired XCI skewing with age is a distinctive feature of blood-derived tissues.

**Fig. 4:**
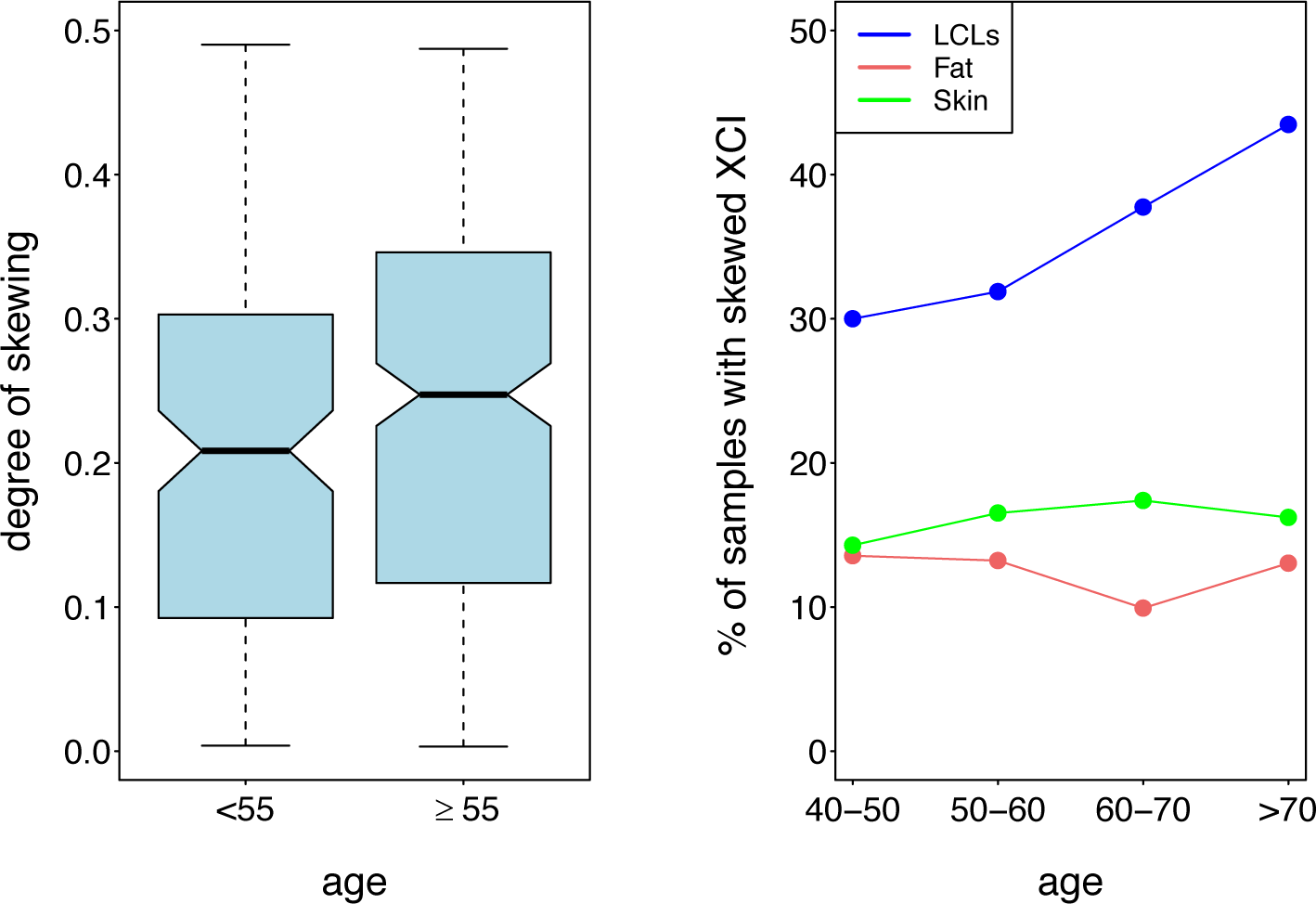
Skewed X-chromosome inactivation (XCI) increases with age in blood-derived tissues but not in fat or skin tissues. Left panel, *boxplots* show the degree of skewing in LCLs samples <55 and ≥ 55. Right panel, *line plots* show the frequencies of skewed XCI at different ages in LCLs (blue), fat (red) and skin (green) tissues.

### Heritability of skewed XCI is dependent on tissue and age

Twin studies are a powerful strategy to investigate the heritability of complex traits. Previous twin studies have reported that skewed XCI in blood-derived samples is heritable, with h^2^ estimates of 0.68 in granulocytes of elderly twin pairs and 0.58 in peripheral blood cells^21-23^, however these studies have not investigated heritability outside of blood. To estimate the influence of additive genetic effects (heritability) and environmental factors on the observed variance in XCI in the three tissues, we implemented the ACE twin model. The ACE statistical model quantifies the contribution of additive genetic effects (A), shared environment (C) and unique environment (E) to the phenotype variance. In order to investigate whether heritability varies with age, we stratified the twin pairs into a younger group (age < 55) and an older group (age ≥ 55; Table S1). Age 55 was chosen as it was identified as the inflection point at which XCI skew begins to increase in the lowess analysis above. We found that XCI skewing is heritable in LCLs of older females (h^2^ = 0.34, *P* = 9.6e-07), but not younger females (h^2^ = 0, *P* = 1). There was no evidence of heritability of XCI skew in fat or in skin tissues at any age (Table 4). The highest proportion of variance was explained by unique environmental factors in all tissues of both younger and older females (E^2^_LCLs_younger_ = 0.99, E^2^_LCLs_older_ = 0.66, E^2^_Fat_younger_ = 0.73, E^2^_Fat_older_ = 0.92, E^2^_Skin_younger_ = 1, E^2^_Skin_older_ = 1). As a complement to the heritability analysis, we calculated intraclass correlation (IC) of XCI skew within MZ and DZ twin pairs of all ages, and within younger and older MZ and DZ twin pairs (Table 5). IC analyses of twin pairs is often used to demonstrate the existence of genetic effect in smaller sample sizes. The IC of XCI skew within MZ twins pairs was positive and statistically significant (IC_MZ_allAges_ = 0.31, *P* = 0.02). We found significant IC of XCI skew within older MZ twin pairs (IC_MZ_older_ = 0.42, *P* = 0.005), but not within young MZ twin pairs (IC_MZ_younger_ = 0.06, *P* = 0.8). We did not detect significant IC within DZ twin pairs at any age, in agreement with previous study in blood^21^. The higher IC of XCI skew within MZ twin pairs compared with DZ twin pairs indicates the involvement of genetic determinants in the regulation of XCI skew in blood-derived tissues. The increase of IC in older compared to younger MZ twin pairs and the fact that the heritability of XCI skew is observed only in females older than 55, confirm a role for genetic variants as age-dependent regulators of the acquired XCI skew in blood-derived tissues. Presumably, genetically-determined secondary cell selection processes act in haematopoietic cell lineages, with the high mitotic rates contributing to the manifestation of their effects in blood-derived tissues. Results also highlight an age-independent role for environmental factors as regulators of XCI skew in blood, fat and skin tissues.

**Table 4.** XCI-skew is heritable is blood-derived tissues of older females. Estimates of the relative contribution of additive genetics, common environment and unique environmental factors to the tissue-specific XCI-skew in age-stratified twins.

| **Tissue [age group]** | **Additive**<br>**genetics (h<sup>2</sup>)** | **Common**<br>**Environment** | **Unique**<br>**Environment** |
| --- | --- | --- | --- |
| LCLs [<55] | 0 | 0.01 | 0.99 |
| LCLs [≥55] | 0.34 | 0 | 0.66 |
| Fat [<55] | 0 | 0.27 | 0.73 |
| Fat [≥55] | 0 | 0.08 | 0.92 |
| Skin [<55] | 0 | 0 | 1 |
| Skin [≥55] | 0 | 0 | 1 |

**Table 5.** Intra-class correlations of XCI skew in age stratified twin pairs. Twin pairs < 55 years-old are classified as young, twin pairs ≥ 55 years-old are classified as old.

|  | **MZ<sub>allAges</sub>** | **MZ<sub>young</sub>** | **MZ<sub>older</sub>** | **DZ<sub>allAges</sub>** | **DZ<sub>young</sub>** | **DZ<sub>older</sub>** |
| --- | --- | --- | --- | --- | --- | --- |
| **n. pairs** | 61 | 18 | 43 | 63 | 25 | 38 |
| **Intra-class**<br>**correlation** | 0.31 | 0.06 | 0.42 | 0 | 0.1 | -0.1 |
| **P-value** | 0.02 | 0.8 | 0.005 | 0.9 | 0.6 | 0.7 |

### XCI skew is associated with smoking status in older females

Tobacco smoking has been reported to induce epigenomic changes including DNA methylation variation (reviewed in^41^). Smoking is a well characterized risk factor in cancer^42^ and, as more recently discovered, in the aetiology of autoimmunity^43^. Although smoking-related X-linked DNA methylation sites have been discovered^44^, no previous studies, to our knowledge, have investigated the relationship between smoking and XCI patterns. We reasoned that changes of XCI patterns may result from smoking, and affect in turn short-term and long-term health. In order to test our hypothesis, we used the 270 individuals in our dataset for which we had smoking status at the time of sample collection, including 233 never smokers and 37 current smokers^45^. We found no difference in the frequency of skewed XCI patterns between never and current smokers (36% and 35% respectively) in LCLs. To take into account the effects of age on the degree of skewing in blood-derived tissues and to examine the relationship between smoking status and degree of skewing at different ages, we split the dataset into a younger (age < 55) and older group (age ≥ 55; Table S2). While the frequencies of skewed XCI were very similar between young smokers and young never smokers (27% and 28% respectively), we detected a higher prevalence of skewed XCI in older smokers compared with older never smokers (47% and 40% respectively). Accordingly, we found an overall positive association between XCI skew and smoking status in older (*P* = 0.02), but not in younger individuals (*P* = 0.5). The data suggest a role for smoking as a modulator of XCI skew in blood-derived tissues of females older than 55. Presumably, the association between smoking and XCI skew changes is complex, and further investigations are needed to characterize the genetic and molecular mechanisms underlying this phenomenon.

## Discussion

In this study, we used multi-tissue transcriptomic data from twins to comprehensively characterize XCI patterns in LCLs, whole-blood, fat and skin tissues from a healthy twin cohort. We show XCI patterns to be tissue-specific and that blood-derived tissues exhibited the highest prevalence of skewed XCI and share the highest similarity of XCI patterns. These findings indicate that XCI patterns are partially driven by tissue-specific mechanisms, and that the XCI skew measured in blood is not a reliable proxy for the skew in other tissues. Skewed XCI patterns limited to disease-relevant tissues and cells have been observed in multiple conditions^9,10,13,14,46,47^ but except for several cases of X-linked diseases, their roles in disease aetiology and predisposition remain largely unknown. Our results demonstrate that tissue-specific XCI patterns within an individual is common in this healthy population.

We show that XCI skewing levels in blood tissues increase with age, with an inflection point at around 55, confirming previous reports^17-21,23,35,39,40^. In this study, more than 41% of females >65 years-old demonstrate skewed XCI patterns in blood-derived tissues, indicating that acquired skewed XCI is a highly prevalent phenotype in ageing populations. We show age-related increase in XCI skew is a distinctive feature of blood-derived tissues, with no evidence for an age-related increase in fat or skin. Age-related increase in XCI skew partially explains the higher incidence of skewed XCI in blood than fat and skin tissues. The effects of age-related skewing of XCI on healthy ageing remain largely unknown, but may have a broad impact on the immune system. Hematopoietic stem cells and the immune system continue to develop throughout life. Presumably, imbalanced X-linked immune-related gene expression toward one parental haplotype leads to a reduced molecular diversity, which may translate in a decline of immune repertoire as well as poor sustenance of the immunological memory.

Previous twin studies have reported that XCI patterns in blood have a genetic component^21,23^. To our knowledge, this is the first study to investigate heritability of XCI skewing levels in other tissues. We found that the heritability of XCI skewing level is limited to blood-derived tissues of females >55 years-old (h^2^ = 0.34), with no evidence of heritability in fat or skin or younger individuals in any tissue. The restriction of heritability to blood of older individuals is of interest given the link between skewed X-inactivation and clonal haematopoiesis. Positive selection of cells carrying an advantageous somatic mutation will lead to clonal haematopoiesis and skewed XCI patterns as the selected cells will carry the same inactivated parental X. Somatic mutation-driven clonal haematopoiesis is now known to be common in blood of healthy older individuals and is often referred to as clonal haematopoiesis of indeterminate potential (CHIP) ^31,48-50^. CHIP is associated with increased risk of both cancer and all-cause mortality^51,52^. The increase in XCI skew in older smokers in our study is consistent with the increase in clonal haematopoiesis observed in smokers^50,53,54^. It is unknown to what extent CHIP accounts for age-acquired XCI skew, however if it is a major driver this would suggest that like age-related XCI skew, CHIP has a significant germline genetic component. Stochastic selection of cells could also contribute to the variance of XCI skewing levels, but, in agreement with previous works ^21,23^, we reason that their contribution is minimal. If stochastic selection of cells was a dominant mechanism, the correlation of XCI patterns between twin pairs would decrease with age.

Overall, the data presented in this study indicate a Gene x Age interaction that shifts the functional allelic dosages of chromosome X in a tissue-restricted manner. The high prevalence of skewed XCI and tissue-restricted XCI in a healthy population could complicate discovery of Chromosome X variants associated with a trait and subsequent genetic risk prediction, as an individual’s genotype may not match their functional genotypic dosage in the relevant tissue. Further investigations of the heterogeneity of XCI patterns across tissues and how this is regulated are essential to clarify the biomedical implications of skewed XCI and its role in healthy ageing in women.

## Materials and methods

### Sample collection

The study included 856 female twins from the TwinsUK registry^28,29^ who participated in the MuTHER study^55^. Study participants included both monozygotic (MZ) and dizygotic (DZ) twins, aged 38-85 years old (median age = 60; Suppl. Fig 1) and were of European ancestry. Volunteers received detailed information regarding all aspects of the research project and gave a prior signed consent to participate in the study. Peripheral blood samples were collected and lymphoblastoid cell lines (LCLs) were generated via Epstein-Barr virus (EBV) mediated transformation of the B-lymphocyte fraction. Punch biopsies of subcutaneous adipose tissue were taken from a photo-protected area adjacent and inferior to the umbilicus. Skin samples were obtained by dissection from the punch biopsies. Adipose and skin samples were weighed and frozen in liquid nitrogen. This project was approved by the research ethics committee at St Thomas’ Hospital London, where all the TwinsUK biopsies were carried out. Volunteers gave informed consent and signed an approved consent form prior to the biopsy procedure. Volunteers were supplied with an appropriate detailed information sheet regarding the research project and biopsy procedure by post prior to attending for the biopsy.

### Genotyping and phasing

Whole-Genome Sequence data (WGS) were generated within the UK10K project as previously described^56^. 557 individuals had both X-chromosome sequence data and RNA-seq data for at least one tissue. For individuals with unavailable X chromosome sequence data, X-linked genotypes data were retrieved from the TwinsUK genotypes previously imputed into the 1000 Genomes Project phase 1 reference panel^57,58^, as described^59^. Haplotypes of X-linked SNPs with a MAF >5% were then phased using shapeit v2.r837^60,61^, with the --chrX flag to set up all functionalities for the phasing of non pseudo-autosomal (non-PAR) regions of the X-chromosome, along with a phasing window of 2Mb and 1000 conditional states. Phasing was performed using the genetic map b37 and the 1000 Genomes Project phase 3 reference panel of non-PAR X-linked haplotypes ^61^. Phased X-linked SNPs were used for further analysis.

### RNA-sequencing data

RNA-sequencing data were generated as previously described^29^. Briefly, the Illumina TruSeq sample preparation protocol was used to generate the cDNA libraries for sequencing. Samples were then sequenced on an Illumina HiSeq 2000 machine and 49 bp paired-end reads were generated. Sequencing reads were aligned to the UCSC GRCh37/hg19 reference genome with the Burrows-Wheeler Aligner v.0.5.9^62^. Genes were annotated using the GENCODE v10 reference panel^63^.

### Longitudinal RNA-sequencing data

Peripheral blood samples were collected 1 to 2.7 years apart from 114 female twins of the TwinsUK registry and were processed with the Illumina TruSeq protocol, sequenced on a HiSeq 2000 machine and 49 bp paired-end reads aligned as described in^36^. Adapter and polyA/T nucleotide sequences were trimmed using trim_galore and PrinSeq tools^64^, respectively. Reads were aligned to the UCSC GRCh37/hg19 reference genome with the STAR v.2.5.2a aligner^65^. Alignments containing non-canonical and unannotated splice junctions were discarded. Properly paired and uniquely mapped reads with a MAPQ of 255 were retained for further analysis.

### Purified immune cells RNA-sequencing data

Monocytes, B, T-CD4^+^, T-CD8^+^ and NK cells were purified using fluorescence activated cell sorting (FACS) from two monozygotic twins exhibiting skewed XCI patterns in LCLs and from 1 individual exhibiting random XCI patterns. Total RNA was isolated and cDNA libraries for sequencing were generated using the Sureselect sample preparation protocol. Samples were then sequenced in triplicates on an Illumina HiSeq machine and 126 bp paired-end reads were generated. Adapter and polyA/T nucleotide sequences were trimmed using trim_galore and PrinSeq tools^64^ respectively. Human and prokaryotic rRNAs were identified using sortmerna v.2.1^66^ and removed. Reads were aligned to the UCSC GRCh37/hg19 reference genome using STAR v.2.5.2a^65^. Alignments containing non-canonical and unannotated splice junctions were discarded. Properly paired and uniquely mapped reads with a MAPQ of 255 were retained for further analysis.

### Correction of RNA-seq mapping biases

To eliminate mapping biases, all RNA-seq data were re-aligned within the WASP pipeline for mappability filtering^67^. The WASP tool has an algorithm specifically designed to identify and correct mapping biases in RNA-seq data. In each read overlapping a heterozygous SNP, the allele is flipped to the SNP’s other allele (generating all possible allelic combinations) and the read is remapped. Reads that did not remap to the same genomic location indicate mapping bias and were discarded. Reads overlapping insertions and deletions were also discarded. Properly paired and uniquely mapped reads were retained for analysis.

### Quantification of allelic read counts and Allele Specific Expression

Allelic read counts at heterozygous SNPs within *XIST* were quantified from paired RNA-seq and X-linked genotypes data with GATK ASEReadCounter^68^. Reads flagged by ASEReadCounter as having low base quality were discarded. To increase the confidence that genotypes were truly heterozygous, only X-linked SNPs with both alleles detected in RNA-seq data and with a read depth of least 10 reads were retained for analysis. X-chromosome allelic read count tables of samples with at least 1 *XIST*-linked SNP passing all quality filters were retained as informative of XCI skewing levels. To quantify the allele specific expression (ASE) of each heterozygous X-linked SNP, the read count at the major allele was divided by the read depth at the site.

### Quantification of *XIST*_ASE_ and degree of skewing of XCI (DS)

In each sample, the XCI skewing levels were quantified by averaging the ASE values of heterozygous SNPs within *XIST*. All SNPs were phased prior to averaging as detailed above. The measure, called *XIST*_ASE_ is defined as follow:

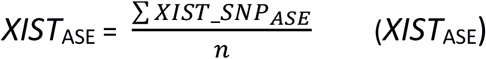

where *XIST*_SNP_ASE_ are the ASE values of heterozygous SNPs within *XIST* and *n* is the number of heterozygous SNPs within *XIST* in the sample. *XIST*_ASE_ is a proxy of XCI skewing levels in a sample, with values ranging from 0 to 1. An *XIST*_ASE_ value of 0.5 indicates equal inactivation of the two parental chromosomes, whereas a value of 0 or 1 indicates complete inactivation of one parental chromosome.

To be consistent with previous literature^21,33^, we classified samples with *XIST*_ASE_ ≤ 0.2 or *XIST*_ASE_ ≥ 0.8 to have skewed XCI patterns, and samples with 0.2< *XIST*_ASE_ < 0.8 to have random XCI patterns. To have an absolute measure of the magnitude of the XCI skewing levels in each sample, (or effect size of *XIST*_ASE_), the degree of skewing of XCI (DS) was calculated. DS is the absolute deviation of *XIST*_ASE_ from 0.5. In each sample, DS was calculated as follow:

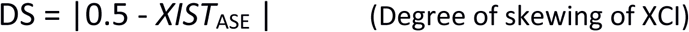

DS does not take into account the direction of XCI skewing, but the degree of deviation from a 50% XCI patterns (*XIST*_ASE_ = 0.5). DS is then a measure of the magnitude of XCI skewing levels in a sample. DS values range from 0 to 0.5, where 0 means random XCI and 0.5 completely skewed XCI patterns. Samples with DS ≥ 0.3 were classified to have skewed XCI, while samples with DS < 0.3 were classified to have random XCI.

### Heritability analysis of DS

The relative contributions of additive genetic factors (A), shared (C) and unique environmental factors (E) to the tissue-specific variance of DS, were calculated using the *twinlm*() function in the *mets* R package^69^. For each tissue, samples were split into a young (< 55) and an older (≥ 55) group according to their ages (Table S1). Due to the low number of MZ and DZ twin pairs in each group, whole-blood was excluded from heritability analysis. To further assess the contribution of genetic effects, the intraclass spearman’s correlation (IC) of DS in blood-derived tissues of young and older MZ and DZ twin pairs was also calculated.

### Association between degree of skewing and smoking status

Association between the degree of skewing in LCLs and self-reported smoking status was tested in the 270 individuals with reliable smoking status recorded^45^. Dataset included 270 females classified either as current smokers (N = 37) or never smokers (N = 233; Table S2). To examine the association between DS and smoking status, the smoking status was converted into a binary trait (0 = no smoker, 1 = smoker). A linear model of the DS as a function of the smoking status was then implemented for younger (age < 55) and older (age ≥ 55) individuals separately. Age was used as covariate. A *P* value ≤ 0.05 was considered to be statistically significant.

## Acknowledgements

This study was supported by MRC Project Grant (MR/R023131/1) to K.S.S. The TwinsUK study was funded by the Wellcome Trust and European Community’s Seventh Framework Programme (FP7/2007-2013). The TwinsUK study also receives support from the National Institute for Health Research (NIHR)-funded BioResource, Clinical Research Facility and Biomedical Research Centre based at Guy’s and St Thomas’ NHS Foundation Trust in partnership with King’s College London. This project was enabled through access to the MRC eMedLab Medical Bioinformatics infrastructure, supported by the Medical Research Council [grant number MR/L016311/1]

## Authors contributions

A.Z. M.D. and K.S.S conceived and designed the project. A.Z performed analysis. SN contributed data discussion. P.C.T and J.T.B. contributed data. SR and C.Y.W performed HUMARA experiments. A.Z. and K.S.S. wrote the manuscript. All authors read and approved the manuscript. The authors also thank Julia El-Sayed Moustafa and Amy Roberts for providing feedback on the manuscript.

## Materials and Correspondence

Correspondence should be addressed to Kerrin S. Small

## Data Availability

TwinsUK RNAseq data is available from EGA (Accession number: EGAS00001000805). TwinsUK genotypes are available upon application to TwinsUK (www.twinsuk.ac.uk).

