## Supplementary Data for "Heritability of skewed X-inactivation in female twins is tissue-specific and dependent on age"

### Supplementary figures

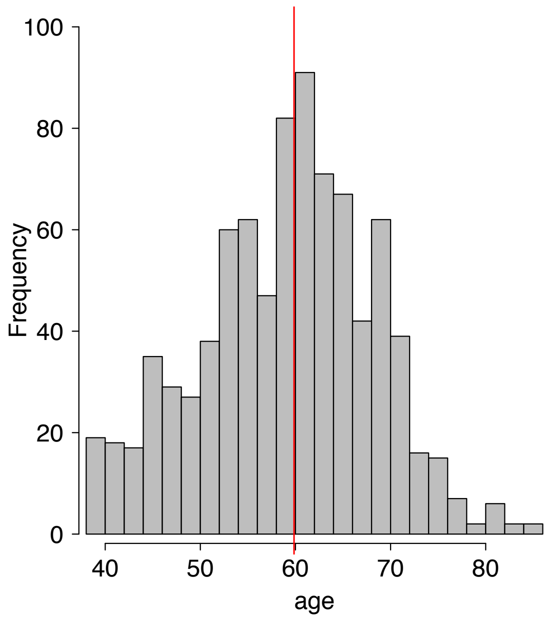


**Fig S1:** Distribution of ages in the TwinsUK samples used in this study. Red line represents the median age.

### Supplementary tables

| Tissue [age group] | Monozygotic twin pairs | Dizygotic twin pairs | Total twin pairs |
|---|---|---|---|
| LCLs [<55] | 18 | 25 | 43 |
| LCLs [≥55] | 43 | 38 | 81 |
| Whole-blood [<55] | 4 | 4 | 8 |
| Whole-blood [≥55] | 7 | 7 | 14 |
| Fat [<55] | 17 | 19 | 36 |
| Fat [≥55] | 34 | 45 | 79 |
| Skin [<55] | 17 | 17 | 34 |
| Skin [≥55] | 27 | 32 | 59 |

**Table S1. Number of monozygotic and dizygotic twin pairs with informative XCI calls for both co-twins in each tissue.**

| | Number of never smokers | Number of current Smokers | Total number |
|---|---|---|---|
| **All ages** | 233 | 37 | 270 |
| **< 55** | 76 | 22 | 98 |
| $\geq$ **55** | 157 | 15 | 172 |

**Table S2. Sample size for smoking analysis.** Younger (age < 55) and older (age $\geq$ 55) women were classified as never smokers or current smokers according to consistency in self-reported questionnaire data taken both at the time of sampling and 5.1 $\pm$ 0.70 years later[45]. Past smokers were excluded from analysis.
